# A retrospective assessment of temperature trends in northern Europe reveals a deep impact on the life cycle of *Ixodes ricinus* (Acarina: Ixodidae)

**DOI:** 10.1101/467811

**Authors:** Agustin Estrada-Peña

## Abstract

This study modelled the changes in the development processes of the health-threatening tick *Ixodes acinus* in northern Europe as driven by the trends of temperature (1950-2016). I used the ECA&D dataset of temperature interpolated at a resolution of 0.25^<u>o</u>^ as the base data for further calculations, which were based on a previously developed process-driven model of the tick. I used the annual accumulated temperature in the period 1950-2016 to obtain the development rates of the oviposition, incubation, larva-nymph, and nymph-adult molts. Annual values were used to ascertain the trend in development rates of each stage. The ecological division of northern Europe (LANMAP2) was used to summarize results along large regions. The temperature in the years 1950-2016 clearly increased in every area of the target territory. The largest increase was observed for a wide territory eastern to Baltic countries, north-eastern Sweden and northern Finland. The development rates of every tested life cycle process had a trend to being faster throughout the time series. Moderate to high increase of the oviposition rates (70%-100% faster) resulted in central Sweden, Baltic countries, parts of Finland, and adjacent territories of Russia. Faster (70%-90%) incubation and molting rates were consistently observed in the same territories and also in large areas of western Norway. The trend of temperature in the period 1950-2016 shows a consistent inflection point around the year 1990, when the slope of the time series of temperature drastically rose. A comparison between 1950-1990 and 1991-2016 demonstrated that annual accumulated temperature was 86% and 26% higher in the Alpine regions, 7%-8% in the Atlantic and 157%, 10% and 16% in Boreal, Continental, and Nemoral regions, respectively. It is concluded that (i) accumulated annual temperature is clearly increasing in the studied territory, (ii) changes were larger since approximately the year 1990, and (iii) these changes have a deep impact on the life cycle of the tick *I. ricinus.* Faster development rates could be part of the processes driving the reported spread of the tick in the target area and should be considered as a serious thread to human health.

## 2. Introduction

World scientists recently released a “Second Warning to Humanity” (Ripple et al. 2017) based on the manifesto issued 25 years ago by the Union of Concerned Scientists (1992). In that manifesto, Ripple et al. (2017) promoted a “call” about the losses of natural landscapes, the progressive deterioration of the landscape, the exhaustion of resources, and, in general, a scientific view of the effects of human kind on climate and other natural resources. It is widely accepted that climate tends to be warmer, with shorter autumns and winters, and with more unpredictable rainfall patterns, impacting the life cycles of many organisms. While studies mostly addressed the impact of the climate trends on free living organisms, there is a lack of information regarding the impact on the life cycle processes regulating the development of ticks of potential importance for human health, because its role in the transmission of pathogens to humans.

These studies were addressed the predicted impact of climate trends on species of plants or animals, commonly using mechanistic models matching known distributions with explanatory variables of diverse quality (Araújo et al., 2005; Parmesan and Yohe, 2003; Pearson and Dawson, 2003). Some of these studies explicitly addressed the expected effects of the climate trends on the distribution and phenology of parasitic arthropods (Hales et al., 2002; Reiter, 2001). This is a topic of interest since many species of Insecta and Acarina (i.e. ticks) are known to be effective vectors of pathogens to humans. Some studies focused on the modelling of each life process of mosquitoes because their importance as vectors of human-affecting pathogens (Lambrechts et al., 2011; Rochlin et al., 2013). The interest on ticks has increased in the last years, mainly after the realization that some human pathogens are (re)emerging, like the agents of Lyme borreliosis (Hanincová et al., 2006; Kimberlin et al., 2015) or the recurrent epidemics and apparent spread of Crimean-Congo hemorrhagic fever (Leblebicioglu et al., 2016; Al-Abri et al., 2017).

Ixodid ticks are ectothermic organisms with a complex life cycle, in which as many as three different stages quest in the vegetation looking for a new host for feeding. After feeding, the tick detaches and molt within the humid places of the ground. Therefore, a change of temperature immediately implies an impact on the development rates of the tick. A faster development does not immediately imply an increased abundance of ticks, because the presence and abundance of suitable hosts play a role in tick abundance (i.e. Vial et al., 2016). While temperature drives the development rates, relative humidity and water saturation deficit outline the mortality rates of the ticks, either while molting or while questing for a host. It has been demonstrated that cold winters may “reset” the population of ticks, greatly reducing the populations of these arthropods (Dautel et al., 2016; Furness & Furness, 2017). It has been pointed out that shorter winters and their effect on ticks and hosts are part of the chain of events leading to larger contact rates between humans and ticks (Randolph, 2004; Sumilo et al., 2007). An increased abundance of ticks could be translated into a higher exposure of humans to them and to the pathogens they transmit, if adequate vertebrate reservoirs are also present (Tälleklint and Jaenson, 2014).

The spread of the tick *Ixodes ricinus* towards northern latitudes is a topic of concern in Europe, as is the case in southern Canada with *Ixodes scapularis* (Leighton et al., 2012; Ogden et al., 2006, 2008). The spread of *I. ricinus* and transmitted pathogens in both latitude and altitude has been already documented by field surveys (i.e. Danielova et al., 2006; Lindgren and Jaenson, 2006; Jaenson and Lindgren, 2011; Jaenson et al., 2009; Materna et al., 2005). Since pathogens have been also detected out of the historical range of the tick it was concluded that vertebrate reservoirs are also being affected by the trends of climate, paralleling the tick spread. It has been however pointed out that climate is not the only cause driving the changing distribution patterns of the tick in Europe, since the availability of suitable hosts for feeding females, like wild ungulates or livestock, is also of central importance (Jore et al., 2011; Tagliapetra et al., 2011). Females fed on large ungulates are able to oviposit literally thousands of eggs, resulting in large populations of larvae the next year, which will contribute to an increased density of the tick. It has been thus suggested that the (re)introduction of wild ungulates at areas of northern Europe may interact with the climate producing the observed changes of the distribution and density of the tick (Jaenson et al., 2012; Medlock et al., 2013).

This study intends to model the effects of the trend of temperature on the development rates of the tick *I. ricinus* in its northern distribution limit. This is the fringe of the distribution of the tick, and thus, effects on the tick’s life cycle would be better observed. I used a long series of interpolated daily values of temperature over Europe for the years 1950-2016 to explicitly evaluate (i) the rate of change of temperature in the period, and (ii) the effects of the changing temperature on the duration of the development periods of the tick. I also evaluated the variability of the impact on the tick’s life cycle in the different ecological regions of Europe aiming for a comparative background along the large geographical divisions of the target region. Results are expected to provide an adequate framework for both adaptation and mitigation in human health.

## 3. Material and methods

### 3.1. Purpose

I aimed to capture the effects of the temperature in the period 1950-2016 on the temperature-dependent processes of the life cycle of the tick *Ixodes ricinus.* The date is the oldest for which available daily climate data covering the European territory exist at an adequate spatial resolution (0.25°). This study is not intended to evaluate the “abundance” or “density” of ticks in a territory, or to calculate the complete life cycle of *I. ricinus* in the target area. No humidity data are available at this resolution, and it is known that water stress in ticks is reduced by the water saturation deficit (**Perret** et al.). Therefore, mortality could not be calculated. In the same way, the length of the questing periods and the implicit mortality in the period could not be calculated because it depends on the density of available hosts and the water stress, a feature that has a local nature. The scarcest the host, the longest the time of questing, and then the highest the water stress. The lack of both data, hots density and water saturation deficit would make unreliable the evaluation of the complete cycle, including peaks of activity or recruitment of stages into the vegetation. Therefore, this study is restricted to (i) evaluate how the temperature-dependent life cycle stages of the tick have been evolving (faster, slower) in a period of 70 years, and (ii) to model how these changes evidenced in the different ecological regions of the target territory. The study is restricted to the northern portions of Europe, where (i) the largest impact of climate should be expected, and (ii) the water stress should have a negligible effect on the ticks since relative humidity and saturation deficit are suitable for the tick (Ogden, pers. comm.), thus increasing the reliability of estimations.

### 3.2. Data on climate and ecological regions in Europe

The climate data were obtained from ECA&D (acronym for European Climate Assessment & Data), which is a dataset of in-situ meteorological observations within Europe gridded into a geographic projection (currently at https://icdc.cen.uni-hamburg.de/1/daten/atmosphere/ecad/ accessed, September 2017). Original climate data include temperature, rainfall and pressure at sea level, at a daily time resolution and 0.25^<u>o</u>^ of spatial resolution (Van den Baselaar et al., 2002). The list of climate recording stations used for interpolations is provided in the web site. Only average daily temperature data for the period 1950-2016 were used. Data were imported from its original netCDF format into the R programming environment (R Core Team, 2018) for further analyses. The calculation of the development rates of the tick on a daily basis would be unrealistic and difficult to compare in a period of 65 years, for multiple ecological regions. I thus opted for the calculation of the accumulated degrees Celsius in one complete year, as base data for input into the equations calculating the life cycle.

I adhered to the scheme of ecological regions of Europe outlined in LANMAP2 (https://www.wur.nl/en/show/The-European-landscape-map.htm, accessed June 2015). The European Landscape Map, LANMAP2, is a pan-European landscape database at a scale of 1:2,000,000. LANMAP2 covers an area of approximately 11 million km^2^ and is a hierarchical classification with four levels and 350 landscape types at its lowest level (more than 14,000 mapping units with an average size of 774 km^2^). The smallest mapping unit is 11 km and the largest is 739,000 km^2^. This study is restricted to the portions included in the coordinates 66^<u>o</u>^N, 25^<u>o</u>^W (top-left) and 47^<u>o</u>^N, 40^<u>o</u>^E (bottom-right). Figure 1 includes the standard climate definitions of the target territory, to which most of the analyses will be related.

**Figure 1.**
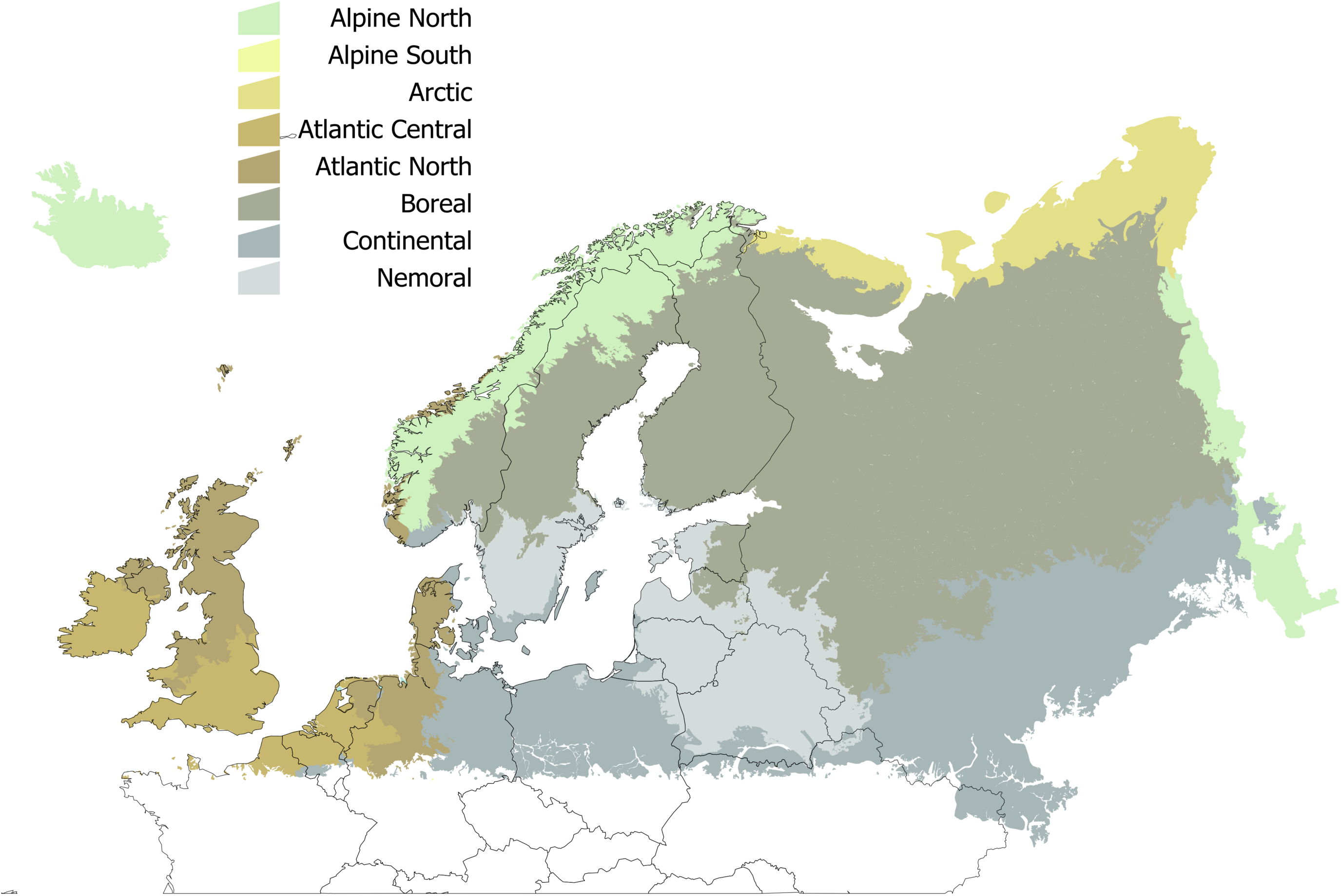
The spatial distribution of the standard definitions of climate (bio-climatic regions) in the target territory, according to the standard denominations of LANMAP2.

### 3.3. Calculation of the development of *Ixodes ricinus*

I aimed to evaluate the annual duration of four development periods of *I. ricinus*, namely (i) the pre-oviposition plus oviposition periods, herein referred as “oviposition”, (ii) the incubation period, (iii) the duration of the larva-nymph molt, and (iv) the duration of the nymph-adult molt, individually for the years 1950-2016. As stated before, I acknowledge that this does not cover the complete life cycle of the tick, but it is aimed to interpret the impact of the temperature on target stages of the life cycle. All the equations used to calculate the development rates of the four stages listed above were previously published together with scripts written in the R programming environment (Estrada-Peña and Estrada-Sánchez, 2013). Therefore, the duration of the oviposition, incubation, larva-nymph, and nymph-adult molts was calculated for every year and for every single 0.25^<u>o</u>^ square for which data exist in the target territory, using the accumulated annual temperature of each single squared cell.

### 3.4. Other calculations

All the calculated data obtained for every single 0.25^<u>o</u>^ cell were transferred into the polygons representing the European climatic regions as outlined in LANMAP2. Each polygon was loaded with the median value of either the accumulated temperature or the development rates, for each year. I calculated the trend of either the accumulated annual temperature or the development rates with functions available in the package ‘greenbrown’ (Forker and Wutzler, 2015) for the R programming environment (R Core Team, 2018). The trend was calculated as the slope of values for the time series between the years 1950-2016, using the function ‘trend’ and the method ‘SeasonalAdjusted’ explicitly addressing an annual seasonal cycle. For every ecological region, I did plot the trends of the accumulated yearly temperature and the development rates (in days) for the period 1950-2016.

I also examined if the series of temperature data had statistically significant break points. A break point is a moment of the complete time series were the data suddenly change its trend. Even if the trend is in the same direction (increasing or decreasing) a break point is an abrupt change in the series, in which data shows an abnormal and sustained behavior different form the one observed in a previous period of time. I examined the break points of every time series (accumulated temperature and the four development rates) using the package ‘changepoint’ (Killick and Eackley, 2014) and the function ‘cpt.meanvar’, available for the R programming environment (R Core Team, 2018). After calculating the existing breakpoints for the temperature series of each single 0.25^<u>o</u>^ cell, I transferred the results into the climatic regions of the target territory for summarizing. I further calculated the values (accumulated temperature, development rates) for the periods of time before and after the breakpoint(s) was/were detected, with an implicit estimation of the difference of values between the two periods of time (pre- and post-breaking point).

## 4. Results

### 4.1. Temperature increased in the target region in the period 1950-2016

The analysis of the annual values of temperature in the time series 1950-2016 demonstrates a clear increase of the temperature in every area of the target territory (Figure 2). The increase of temperature was smaller in areas of Ireland, western and northern United Kingdom and south-western Norway. The greatest increase was observed for a wide territory eastern to Baltic countries, north-eastern Sweden and northern Finland. The large territory in Russia (Figure 2) with a slope of “0” resulted from the lack of interpolated climate data; it remains in the figures because it is part of the LANMAP2 ecological description of the territory. No slope less than 0 has been observed (i.e. a trend to decrease of temperature). Table 1 summarizes data for the seven bioclimatic regions enclosed in the target territory (according to the LANMAP2 scheme). The table 1 also includes the data a break point in the trend of temperature that has been consistently observed around the year 1990. I therefore calculated the mean annual accumulated temperature separately for the periods 1950-1989 and 1990-2016, with an explicit estimation of the percent of the difference. For every region, the mean accumulated annual temperature is always higher in the period 1990-2016, at a variable percent depending on the latitude: northern regions experienced a more abrupt change.

**Figure 2.**
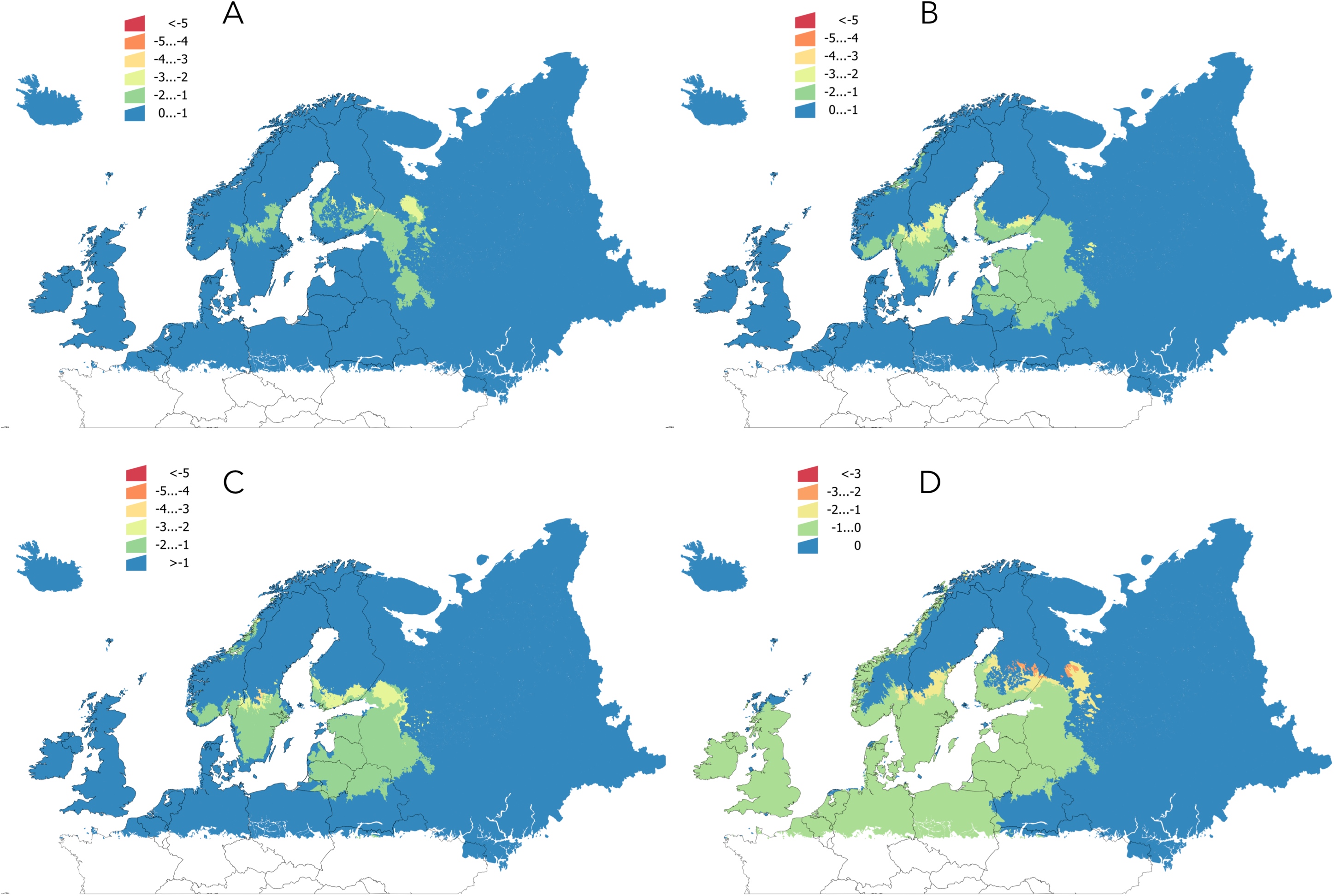
Changes of temperature in the time series 1950-2016. A: the slope of the changes of temperature of the complete period. B: the averaged annual accumulated ^<u>o</u>^C in the period 1950-1989. C: the averaged annual accumulated ^<u>o</u>^C in the period 1990-2016. D: the percent of changes of temperature between the period 1990-2016 and 1950-1989. The large territory in Russia with a homogeneous color are areas where no temperature data are available. Minute polygons of the territory displaying “no change” across Europe are small areas smaller then the resolution of the layers of temperature (the size of the patch is smaller than 0.25^<u>o</u>^ and therefore the spatial transfer of the information from raster to the vectorial layer cannot result in reliable data).

**Table 1.** Values of the annual accumulated temperature (°C) in the target territory, summarized according to the standard climate classification of Europe. Included are the trend for the period 1950-2016, and the annual accumulated temperature, averaged for the periods 1950-1989 and 1990-2016, since a clear inflection point in the trend of temperature was noticed around the year 1990. The “% of change” indicates the difference of the accumulated annual temperature between the time series 1950-1989 and 1990-2016.

| **Climate category** | **Trend**<br>**(1950-2016)** | **Annual**<br>**accumulated temperature:**<br>**1950-1989** | **Annual**<br>**accumulated temperature:**<br>**1990-2016** | **% change between 1950-1989**<br>**and 1990-2016** |
| --- | --- | --- | --- | --- |
| Alpine North | 8.38 | 0.96 | 267.00 | 86.90 |
| Alpine South | 8.91 | -2.19 | 65.32 | 26.29 |
| Atlantic Central | 7.63 | 3492.24 | 3745.04 | 7.25 |
| Atlantic North | 7.17 | 2871.40 | 3107.71 | 8.37 |
| Boreal | 10.66 | 742.51 | 1089.42 | 157.09 |
| Continental | 9.15 | 2890.30 | 3190.26 | 10.49 |
| Nemoral | 10.56 | 2135.75 | 2480.61 | 16.50 |

**Table 2:**
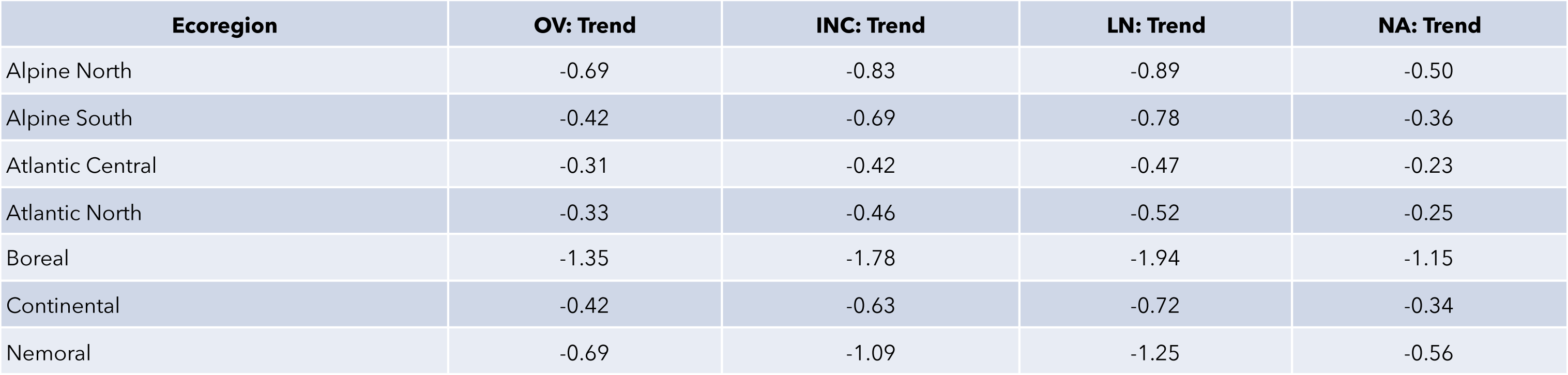
Trends in the four modelled developmental stages of the life cycle of *I. ricinus* grouped according to the ecoregion in Europe (LANMAP2). OV: Oviposition, INC: Incubation, LN: larva to nymph, NA: nymph to adult. The negative trend means for a shortening of processes, the larger the negative number, the shorter the period.

| **Ecoregion** | **OV: Trend** | **INC: Trend** | **LN: Trend** | **NA: Trend** |
| --- | --- | --- | --- | --- |
| Alpine North | -0.69 | -0.83 | -0.89 | -0.50 |
| Alpine South | -0.42 | -0.69 | -0.78 | -0.36 |
| Atlantic Central | -0.31 | -0.42 | -0.47 | -0.23 |
| Atlantic North | -0.33 | -0.46 | -0.52 | -0.25 |
| Boreal | -1.35 | -1.78 | -1.94 | -1.15 |
| Continental | -0.42 | -0.63 | -0.72 | -0.34 |
| Nemoral | -0.69 | -1.09 | -1.25 | -0.56 |

**Table 3:**
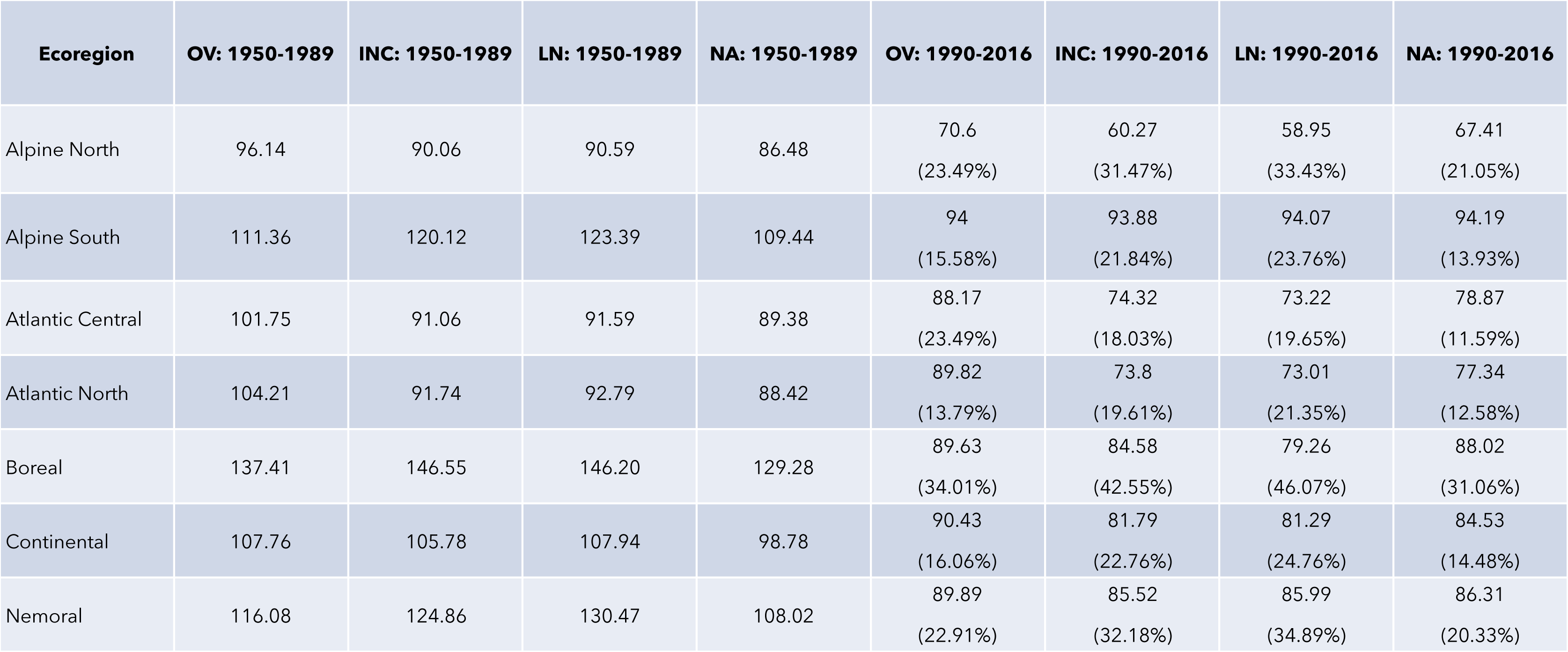
Changes in the four modelled developmental stages of the life cycle of *I. ricinus* grouped according to the ecoregion in Europe. OV: Oviposition, INC: Incubation, LN: larva to nymph molt, NA: nymph to adult molt. The table includes the modelled duration (days) of the stage according to the annual accumulated temperature in the periods 1951-9189 and 1990-2016. The second line in the columns of the period 1990-2016 means for the percent of difference between the time periods.

| **Ecoregion** | **OV: 1950-1989** | **INC: 1950-1989** | **LN: 1950-1989** | **NA: 1950-1989** | **OV: 1990-2016** | **INC: 1990-2016** | **LN: 1990-2016** | **NA: 1990-2016** |
| --- | --- | --- | --- | --- | --- | --- | --- | --- |
| Alpine North | 96.14 | 90.06 | 90.59 | 86.48 | 70.6<br>(23.49%) | 60.27<br>(31.47%) | 58.95<br>(33.43%) | 67.41<br>(21.05%) |
| Alpine South | 111.36 | 120.12 | 123.39 | 109.44 | 94<br>(15.58%) | 93.88<br>(21.84%) | 94.07<br>(23.76%) | 94.19<br>(13.93%) |
| Atlantic Central | 101.75 | 91.06 | 91.59 | 89.38 | 88.17<br>(23.49%) | 74.32<br>(18.03%) | 73.22<br>(19.65%) | 78.87<br>(11.59%) |
| Atlantic North | 104.21 | 91.74 | 92.79 | 88.42 | 89.82<br>(13.79%) | 73.8<br>(19.61%) | 73.01<br>(21.35%) | 77.34<br>(12.58%) |
| Boreal | 137.41 | 146.55 | 146.20 | 129.28 | 89.63<br>(34.01%) | 84.58<br>(42.55%) | 79.26<br>(46.07%) | 88.02<br>(31.06%) |
| Continental | 107.76 | 105.78 | 107.94 | 98.78 | 90.43<br>(16.06%) | 81.79<br>(22.76%) | 81.29<br>(24.76%) | 84.53<br>(14.48%) |
| Nemoral | 116.08 | 124.86 | 130.47 | 108.02 | 89.89<br>(22.91%) | 85.52<br>(32.18%) | 85.99<br>(34.89%) | 86.31<br>(20.33%) |

### 4.2. The development rates of *I. ricinus* life cycle are faster

The values of the development rates of every tested life cycle process of *I. ricinus* had a negative slope in the whole target territory, meaning for a trend to being faster throughout the time series. No regions with a positive slope (i.e. a decrease of development rates) were found. However, the different life cycle stages were not affected in the same magnitude by changes of climate (Figure 3). Moderate to high increase in oviposition rates were found in central Sweden, parts of Finland, and adjacent territories of Russia. The impact of climate on the incubation rates is more noticeable. Areas of a faster incubation extend over large regions of central and southern Sweden, southern Finland, most of Baltic countries and adjacent territories of Russia for which data are available. The slope of changes of the larva-nymph molt rates has a similar geographical distribution, but the effects of the climate on this stage extend far southern Sweden and are more noticeable in southern Finland and adjacent Russian territories. A faster nymph-adult molt has been observed in most of the explored territory. The impact is maximum in central Sweden and Finland (large negative slope). It is the only detectable life cycle process that had obvious changes in the complete period of time (1950-2016) in coastal Norway.

**Figure 3.**
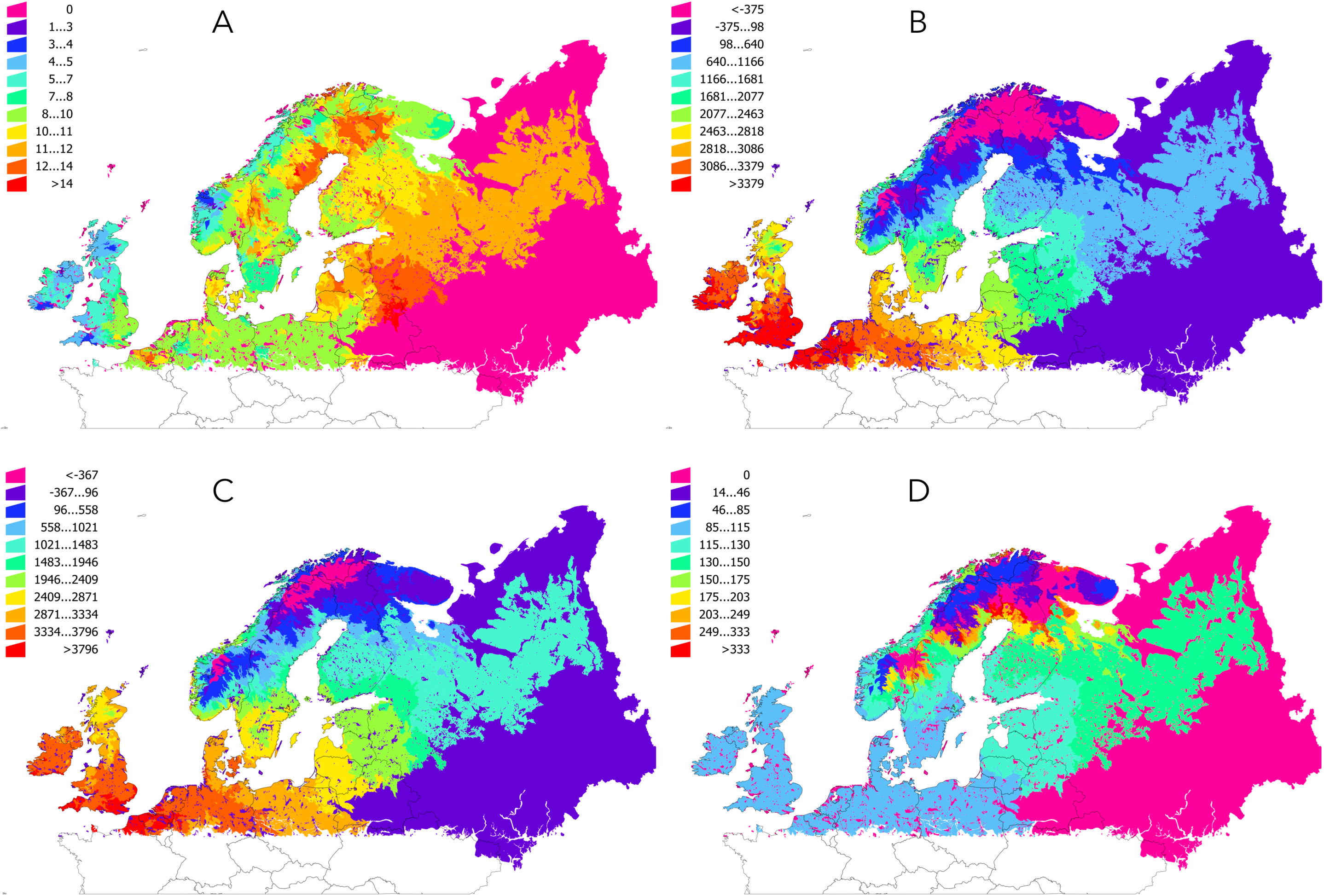
Trends of the development rates of four processes of the life cycle of *Ixodes ricinus.* A: oviposition; B: incubation; C: larva-nymph molt; D: nymph-adult molt. A negative trend means for faster development rates.

The slope of the development rates gives only a general overview of the trend in a long time period. Since I detected an obvious and abrupt change of slope between the years 1950-1989 and 1990-2016, I calculated the differences in the development rates averaged for both periods of time (in %). The results are summarized in Figure 4, in which data are shown in percent of change (i.e. a change of 100% means for a double development rate or, in other words, for half the time in process development). The results display rates of change of about 74%-90% faster in large areas of central Europe, United Kingdom, Ireland, coastal Norway, southern and central Sweden, southern Finland and the Baltic countries.

**Figure 4.**
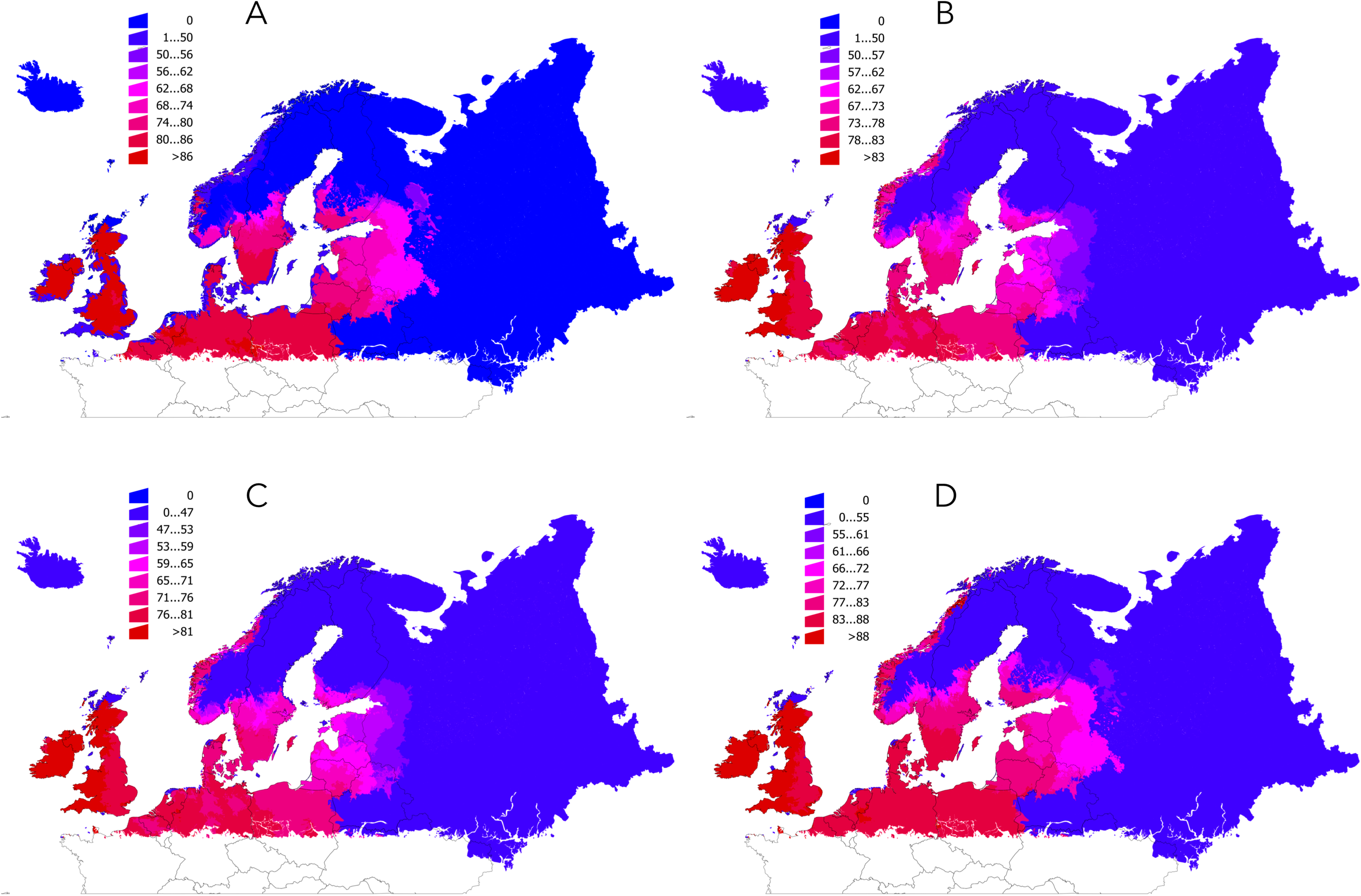
The percent of change of the development rates of the four processes studied for *Ixodes acinus* between the period 1950-1989 and 1990-2016. The data are shown in percent of change, i.e. a change of 100% means for twice the speed of development in the period 1990-2016 in comparison with the period 1950-1989. Development rates for each period were produced with the averaged annual accumulated temperature for both periods of time. A: oviposition; B: incubation; C: larva-nymph molt; D: nymph-adult molt.

The impact of the trends of temperature on the development rates of *I. ricinus* is not homogeneous along a latitudinal range of the target region because the different rates of change of temperature. Figure 5 displays the percent of difference in these rates (higher values mean for faster development). Highest changes in development rates were noticed northern to 57^<u>o</u>^-58^<u>o</u>^N, meaning for a deeper effect of the trends of climate on development rates of the tick. The less affected process is the oviposition. Nevertheless, both molts (larva to nymph and nymph to adult) are the most impacted above the latitudinal limit of 57^<u>o</u>^-58^<u>o</u>^N, with rates of development up to 30% to 68% faster for these two stages.

**Figure 5.**
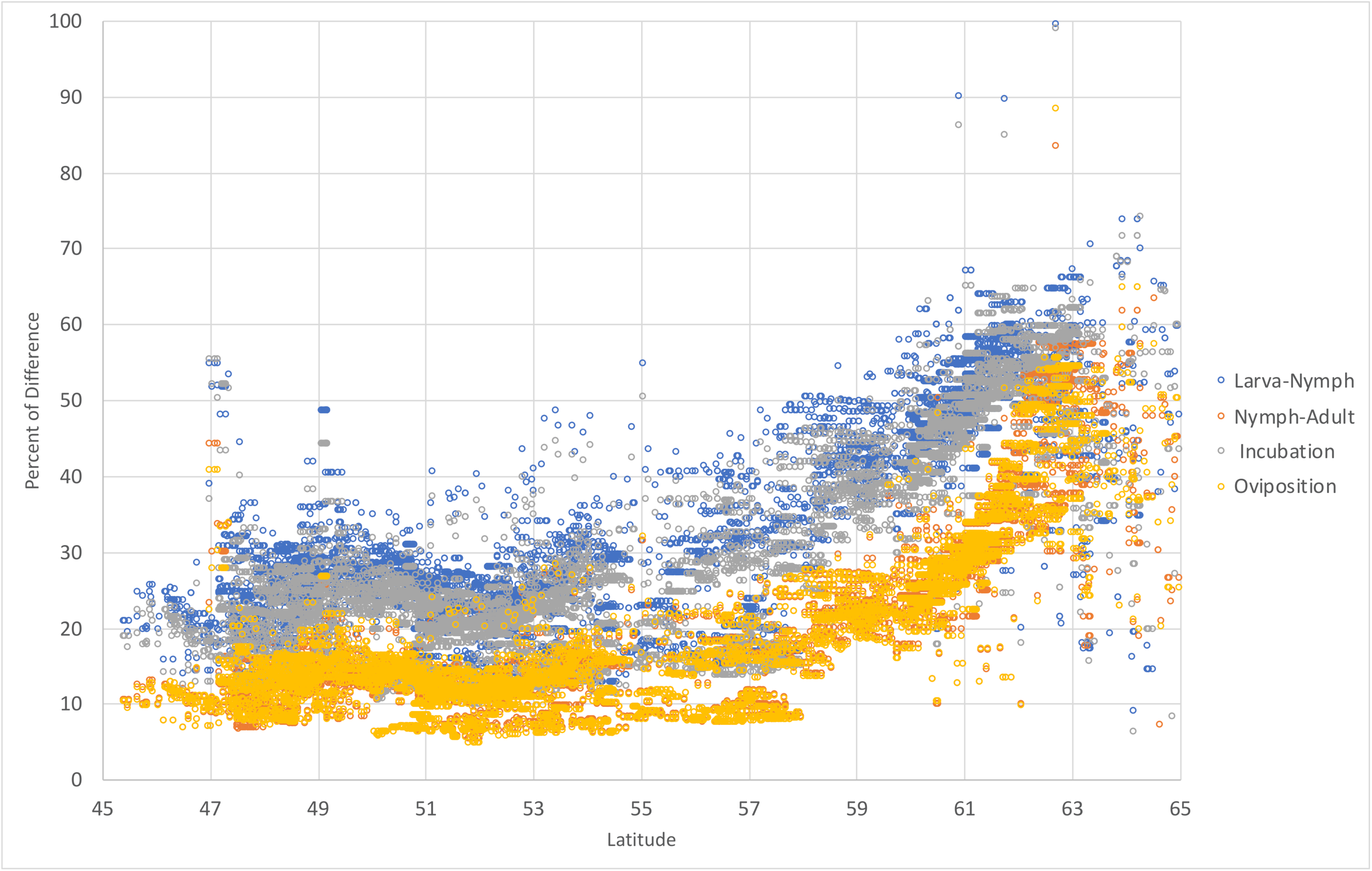
Modelled percent of change of the development rates of the four processes studied for *Ixodes ricinus*, in the range 45^<u>o</u>^N-65^<u>o</u>^N in nordic European countries. The chart displays the percent of difference, computed as the percent between the rates of development in the period 1950-1989 and 1990-2016 (i.e. a value of 100% would represent a development process performed in half of the time). Each point refers to a confirmed record of the tick reported with a pair of coordinates.

## 5. Discussion

This study addressed the effects of the changes of temperature in northern Europe on the development rates of the tick *I. ricinus* in the period 1950-2016. Like any other ectothermic organisms, ticks are deeply affected by changes of temperatures. This has been a reason of concern and has been pointed out as one of the factors driving the colonization of the tick at northern latitude in Europe. It is widely agreed that the increase of temperature is one of the factors that could spread the tick into nordic European countries, northern to its historical distribution range. Studies by Jaenson et al. (2012), Jore et al. (2011) and Laaksonen et al. (2017) have all confirmed the spread of the tick in areas where it was absent only 15-20 years ago, according to field surveys at the time. Climate and vegetation impact tick survival because they are determinants of the occurrence of suitable communities of hosts, and a refuge protecting ticks for off-host desiccation. Second, host-seeking activity is affected by ambient temperature and humidity. Third, rates of development of ticks from one life stage to the next depend on temperature, being faster at higher temperature.

I used an interpolated dataset of temperature (ECA&D) for calculating the accumulated annual temperature and evaluating the rates of development of the tick’s processes for each year. The dataset is publicly available and holds adequate daily data of European climate recording stations. One of its advantages is its high resolution, suitable for addressing models covering biological processes. We previously promoted the use of satellite imagery for the evaluation of the life cycle of ticks (i.e. Estrada-Peña et al., 2016) but the relatively short series of data available from different satellites prevents large periods of time to be evaluated. Since this is a retrospective study, I aimed to have a long climate series. Only interpolated climate datasets have the length necessary to capture the general patterns of change in the life processes of living organisms. Other datasets, like CRU (http://www.cru.uea.ac.uk/data) are of coarser resolution and I opted to use the fine grain of the ECA&D dataset.

This study could not use the basic reproduction number, R_0_, which is the universally accepted value translatable to all branches of epidemiology including those involved in studies of arthropod vector dynamics (Wu et al., 2013). This was because the climate dataset lacks data on water vapor. Therefore, the saturation deficit (affecting survival of the ticks) could not be evaluated in equations addressing the mortality of the tick. The focus was paid only on the impact of the temperature on the development rates of *I. ricinus.* I however assumed, as is the case of *I. scapularis* in southern Canada, that the low water deficit in the region is not responsible of large mortality rates of the tick (Ogden et al., 2014).

Previous studies describing the changes of environmental suitability for *I. ricinus* in Europe suggested that temperature is the limiting factor for the establishment of the tick in northern latitudes, while depletion of water vapor in the air is the limiting factor in its southern distribution limit (Estrada-Peña et al., 2006; Jaenson et al., 2012). Previous studies on the topic were based on basic mechanistic models involving several variables of climate (Estrada-Peña et al., 1999). However, after the development of a process-driven model for *I. ricinus* (Estrada-Peña and Estrada-Sánchez, 2014) it resulted obvious than the physiological processes of the tick are the best representation of the environmental limits for the tick’s survival. While mechanistic, algorithm-driven models, can provide an adequate estimation of the distribution range of an organism, the limiting factors shaping its life cycle are better described by equations tailored for describing life cycle processes (Maino et al., 2016).

Time series analysis uncovered a strong signature of change around the year 1990. This is consistent across the complete geographical range examined. The number of climate recording stations used for both periods of time was approximately the same in both periods (pre- and post-1990) and the interpolation procedures were the same. Therefore, since this is not an issue derived from the procedures preparing the dataset, it should be assumed that there is a clear change of trend. Such change points to a faster increase of temperature, noticed in the complete region around the year 1990. This break makes two clear periods of temperature increase, with a small change of slope in the Nemoral, Continental, Atlantic North, and Atlantic Central regions (7-16% higher temperature in 1990-2016 *versus* 1950-1989), largely noticed in the Boreal and Alpine climatic regions (86% to 157% higher temperature comparing both periods of time). The impact of temperature trends are thus different according to the regions of the target area, and these changes are most noticeable in the northern range of the region studied.

Similar studies exist in southern Canada for the prominent tick species *I. scapularis* (Ogden et al., 2006, 2008, 2014; Wu et al., 2013). In southern Canada, the concerns of spread of that tick and the transmitted pathogens are similar to those for nordic countries in Europe and *I. ricinus.* These studies suggested that temperature is the only limiting factor for the establishment of *I. scapularis* in Canada, but that the observed trends of climate would permit or accelerate the spread of the tick in Canada (Githeko et al., 2000; Parmesan and Yohe, 2003). The conclusions emanating from these reports, some of them made on the use of R_0_ as the measuring value of change of the life cycle, are clear and compatible with our results: *I. ricinus* is experiencing a clear spread towards northern latitudes. Such spread is driven not only by increased host availability but by the prevailing climate, which is promoting faster development rates. The relative importance of both climate trends and increased host availability is still unmeasured by field surveys, but I anticipate it will be hard to separate both effects in field studies.

The impact of the trends of temperature is not the same in every part of the large territory analyzed. This seems to be derived from the sensitivity to change of the different parts of the examined territory, and the different tick processes, a concept already explored by Estrada-Peña and Estrada-Sánchez (2013). Areas in which temperature is well into the range of optimum values for a successful tick’s life cycle, are less prone to observe drastic changes. However, in the fringe of the tick distribution, small changes of temperature would promote that sites far from the optimum climate conditions for the species could experience drastic variations in the tick’s life cycle. Data derived from either single records of the tick (i.e. with coordinates) or summarized along large bio-climatic regions of Europe are conclusive: northern Europe is experiencing the largest impact of the climate on the development rates of the tick. These calculations match well the published reports by i.e. Jaenson et al. (2012), Jore et al. (2011) and Laaksonen et al. (2017) regarding the spread of the tick into European nordic countries.

The effects of such changes in the tick’s life cycle cannot be ignored in the context of human health. Faster development rates in an environment in which water deficit is not a significant driver of tick mortality would probably denote higher tick density, in a territory in which large ungulates are common or even re-introduced culminating in an abundant blood source for adult ticks (Tälleklint and Jaenson, 2014; Jaenson et al., 2018). This should result in an increase of cases of tick-transmitted pathogens to humans in the area, at a yet unevaluated rate (i.e. Sormunen et al., 2016). These challenges to public health systems in northern European regions must to be addressed with an adequate preparation for the expected impact.

## 6. Conclusion

Temperature has changed at an unprecedented rate in Europe in the last decades, the slope of increase after the year 1990 being clearly higher than for the period 1950-1989. These trends led to large rates of change of the development process of the tick *I. ricinus* in regions of northern Europe. The application of a process-based model identifying the changes of the tick’s life cycle demonstrated that oviposition, incubation and molting processes would be impacted by the accumulated annual temperature, resulting in faster development rates. While the presence of adequate hosts for tick feeding is a further element affecting the establishment of permanent populations of *I. ricinus*, the changes of temperature affected the progression of the tick population at its northern fringe. This is of indudable concern under the light of the general World’s trends of climate, and has been already pointed out for other arthropod vectors. The joint spread of the tick vector, together with an expected similar event of pathogen’s vertebrate reservoirs will introduce emerging pathogens into areas that were free o them until a few years ago.

## 7. Acknowledgments

This study has been conducted under the initiative “World Scientists’ Warning to Humanity: a Second Notice” led by William J. Ripple (University of Oregon, USA). Most of the records of *I. ricinus* were collected by the author under the ICTTD initiative led by Frans Jongejan, and funded by European Union in the FP7 program. Adicional data were provided by Thomas G.T. Jaenson and his team (University of Uppsala, Sweden) and Tero Klemola and his team (University of Helsinki, Finland). Andrei D. Mihalca (University of Cluj-Napoca, Romania) coordinated the efforts to improve the dataset of the tick *I. ricinus* in Europe under the European COST action TD1202.

